# Mutual inhibition of airway epithelial responses supports viral and fungal co-pathogenesis during coinfection

**DOI:** 10.1101/2022.04.13.488236

**Authors:** Patrick Dancer, Adam Pickard, Wiktoria Potocka, Kayleigh Earle, Rachael Fortune-Grant, Karl Kadler, Margherita Bertuzzi, Sara Gago

## Abstract

Awareness that fungal coinfection complicates viral respiratory infections causing worse disease outcome has recently emerged. The environmental fungus *Aspergillus fumigatus* (*Af*) has been reported as the main driver of fungal coinfection in patients suffering from viral infections caused by Cytomegalovirus, Influenza or more recently SARS-CoV2. The airway epithelium is the first common point of contact between inhaled pathogens and the host. Aberrant airway epithelial cell (AEC) responses against fungal challenge have been described in patients susceptible to aspergillosis. Therefore, it is likely that a dysregulation of AEC responses during fungal-viral coinfection represents a potent driver for the development of fungal disease. Here we used an *in vitro* model of *Af*-viral infection of AECs to determine outcomes of spore internalisation, killing and viral replication during coinfection. Our data indicate that viral stimulation, while boosting *Af* uptake by AECs, limits *Af* spore killing by those cells, favouring fungal persistence and growth. Type I viral-induced interferon release was significantly decreased in the presence of *Af* hyphal forms suggesting a possible role of *Af* secreted factors in modulating viral pathogenicity. We next explored the impact of *Af* challenge in SARS-CoV2 replication within airway epithelial cells using nano-luciferase as a measure of viral replication. We found that *Af* increased SARS-CoV2 pathogenicity in a strain-dependent manner. Collectively, our findings demonstrate a mutual inhibition of antifungal and antiviral AEC responses during *Af-*viral coinfection and also suggest that some fungal factors might be key regulators of co-pathogenicity during in lung infection.

## IMPORTANCE

Severe viral infections have emerged as a new risk factor for the development of pulmonary aspergillosis. As mucosal responses are critical to prevent respiratory infections caused by a variety of inhaled pathogens, we hypothesised that subversion of these responses during coinfection is at the base of the increased susceptibility of patients suffering viral infections to pulmonary aspergillosis. In this study, we discovered that in the presence of a viral mimicker, airway epithelial cells (AECs) are less able to kill *Aspergillus fumigatus (Af)* spores and this correlates with aberrant cytokine responses. Similarly, when AECs were exposed to SARS-CoV we observed *Af* can sustain viral pathogenesis in a strain-dependent manner. Understanding the mechanisms of AECs responses during fungal-viral coinfection will allow to develop new strategies for the clinical management of these emerging coinfections.

### OBSERVATION

Severe bacterial pneumonia complicating Influenza or more recently COVID-19 have been widely linked with worse disease outcome and increased mortality (1–3). However, awareness that fungal coinfections complicate viral respiratory disease has only recently emerged. During the H1N1 pandemic, influenza associated pulmonary aspergillosis doubled mortality risk (4, 5). Similarly, a 30% reduction in survival has been reported in patients with coronavirus associated pulmonary aspergillosis (CAPA) (6, 7). Cytomegalovirus infections are also known to increase aspergillosis risk in transplant recipients (8). Nevertheless, the mechanistic understanding of fungal-viral coinfections is very limited.

Polymicrobial infections are complex and coinfecting pathogens may interact synergistically or antagonistically compared with single infections. Viral coinfections are commonly associated with a process called viral interference, whereby one of the viruses inhibits the replication of the other virus (9). Using an *in vitro* model of differentiated human bronchial epithelial cells it has been shown that human rhinoviruses (HRVs) inhibited SARS-CoV2 replication (10) as previously reported for HRVs and influenza A coinfections (11). However, increased co-pathogenesis occurs during concomitant infections by *Streptococcus pneumoniae, Staphylococcus aureus, Haemophilus influenzae* and/or *Streptococcus pyogenes* with Influenza [for review (12)]. Nevertheless, the outcome of combined infections depends on both the direct interaction between the coinfecting pathogens and the way by which these interactions modulate the host responses.

The airway epithelium is the first point of contact between inhaled pathogens and the host (13, 14). As such, *Af* spores can adhere to and be internalised and killed by AECs, triggering the downstream activation of antifungal responses (15–21). Accordingly, aberrant epithelial cell responses are major contributors to susceptibility to *Af* infections in at risk-patients (16, 22, 23). Furthermore, during viral exposure, AECs are the primary site where respiratory viruses enter, replicate and disseminate in a process leading to the production of immune mediators (24). Oxidative stress during respiratory viral infections is known to drive the disruption of the airway epithelia barrier integrity thus, facilitating infection by other microorganisms (25). However, little is known about the cellular mechanisms by which viral infection underpins susceptibility to aspergillosis. With a view to understand AECs responses during fungal-viral coinfection, we performed *Af*–viral coinfection of immortalised and primary human AECs by combining the use of genetically-engineered red-fluorescent *Af* strains, the viral mimicker FITC-Poly (I:C) and a nano-luciferase tagged replication competent SARS-CoV2 virus. Using imaging flow cytometry (IFC), live-cell fluorescent microscopy and a nano-luciferase activity assay, we quantitatively measured outcomes of the *Af*–viral coinfection in AECs.

### Viral stimulation impairs *Af* killing by AECs

To assess whether concomitant viral–fungal coinfection affects *Af* spore internalisation by cultured airway epithelia, A549 monolayers were incubated for 6 h with *Af* A1160^+/tdT^ in the presence or absence of FITC-polyriboinosinic:polyribocytidylic acid [Poly(I:C)], a stable synthetic dsRNA analogue frequently used as a TLR3 ligand to mimic viral infection (26). As from an established experimental pipeline (23, 27), extracellular *Af* conidia were stained using calcofluor white, and the number of individual epithelial cells which had internalised *Af* spores (AEC_i_) was determined using IFC. In the presence of FITC-Poly(I:C), *Af* spore uptake by AECs increased 18-fold compared to *Af* A1160^+/tdT^ only challenges (P < 0.05) **(Figure 1a)**. Lipofectamine, which was used to deliver Poly(I:C) into the cells, did not influence *Af* internalisation by AECs, nor did the presence of *Af* alter the interaction between AECs and viral mimicker, as measured via mean fluorescence FITC Poly(I:C) **(Supplementary Figure 1a-b)**.

**Figure 1:**
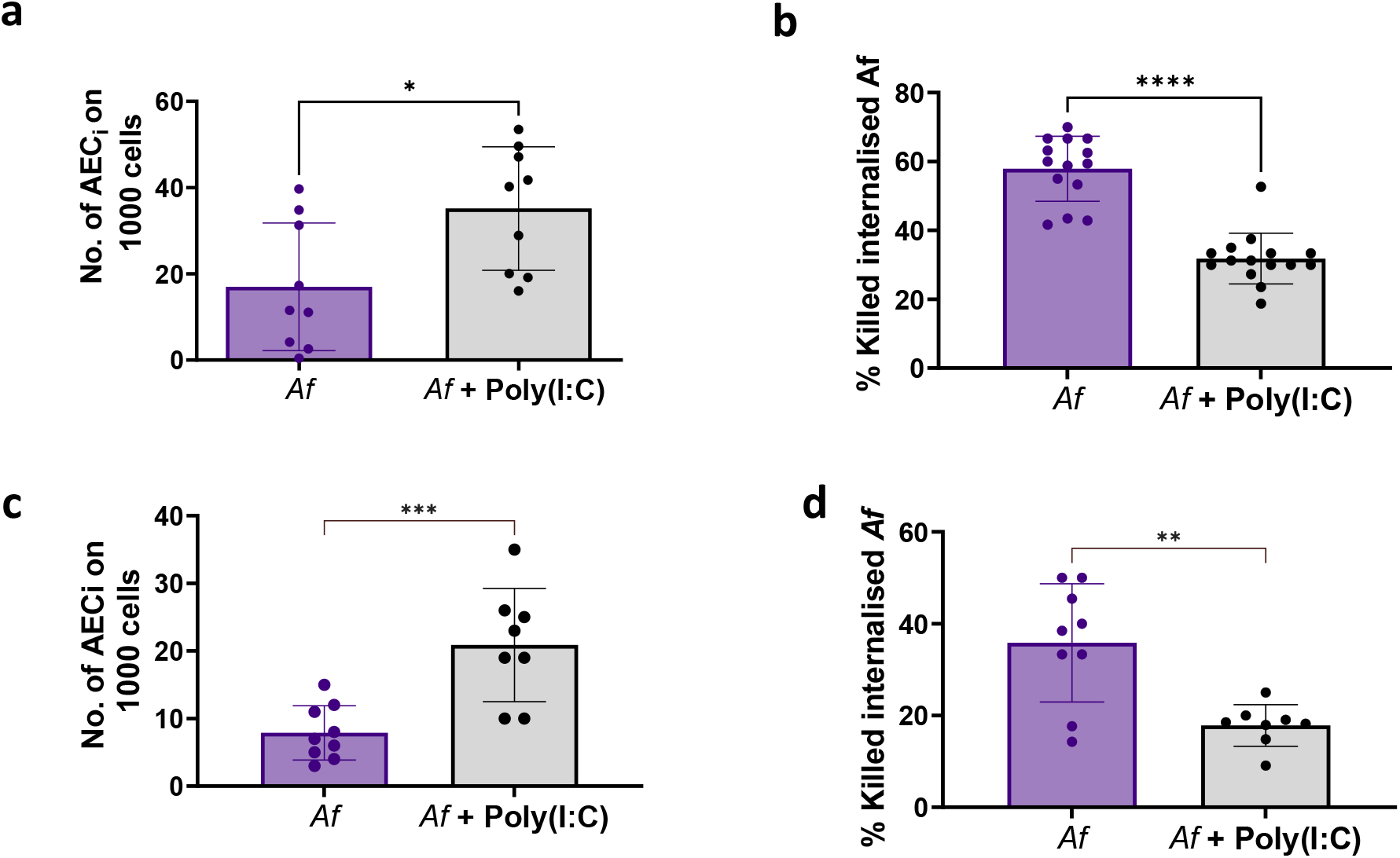
Viral mimic [Poly(I:C)] decreases *A. fumigatus (Af)* intracellular killing by A549 alveolar epithelial cells and primary human AECs. (a) Quantification of AECi on 1000 A549 alveolar epithelial cells by IFC. (b) Percentage of internalised *Af* killed by AECs. (c) Quantification of AECi on 1000 human primary AECs (n = 3 donors) by IFC. (d) Percentage of internalised *Af* killed by human primary AECs (n = 3 donors). Uptake experiments (a,c) were performed at 6 h, while killing experiments (b, d) were performed at 16 h. Data in graphs are represented as mean ± standard deviation of biological and technical triplicates. Mean differences were analysed using either T-test or Man-Whitney test according to normality distribution. \*\*\*\**P* ≤ 0.0001, *** *P* ≤ 0.001, ** *P* ≤ 0.01, and * *P* ≤ 0.05.

*In vitro, in vivo* and *ex vivo* studies by us and others have shown that AECs can kill internalised *Af* spores (15–19, 23). In order to measure *Af* viability, intracellular quenching and lack of germination of A1160^+,mSca^ spores in the presence or absence of FITC-Poly(I:C) was determined using live-cell imaging at 24 h post-infection. Our results indicate that viral stimulation decreases intracellular killing of *Af* spores by 10% compared to single *Af* infection (P<0.05) **(Figure 1b)**. Viability counts of *Af* spores at 8 h post infection also revealed a similar increase of the fungal population in the presence of the viral mimic FITC-Poly (I:C) **(Supplementary Figure 2a)**.

Importantly, the increase in *Af* uptake and decrease of intracellular *Af* killing was corroborated in primary human AECs obtained from 3 healthy donors (D1-3) **(Figure 1c-d)**. Collectively, these findings indicate that viral stimulation, while initially boosting *Af* uptake by AECs, limits *Af* spore killing by AECs, favouring intracellular persistence and likely further damage to the infected monolayers.

### Fungal exposure facilitates viral pathogenesis within AECs in a strain dependent manner

In order to further characterise the outcome of fungal-viral interactions in epithelial cells responses, the release of IL-6, IL-8 and IFNβ by A549 monolayers challenged with *Af* spores in the presence or absence of Poly(I:C) was measured at 9, 16 and 24 h post infection. Our data indicates fungal-viral combined exposure does not impact IL-6 and IL-8 release **(Figure 2a)**. However, *Af* germination was linked to a significant decrease of Poly(I:C) driven IFNβ release (P<0.05) as previously reported for *Alternaria alternata* (28), thus suggesting a possible fungal-driven inhibition of antiviral responses **(Figure 2a)**. Aberrant activation of AECs responses during fungal and viral co-pathogenesis did not result in increased cell death **(Figure 2b)**. On the basis of these observations we further hypothesised that secreted factors produced by *Af* during vegetative growth are likely regulators of viral pathogenicity. To model viral-fungal coinfection during fungal germination, A549 monolayers were exposed to live SARS-CoV expressing a Nanoluciferase reporter (SARS-CoV-2-ΔOrf7a-NLuc (29)) in the presence or absence of *Af* A1160 and Af293 culture filtrates recapitulating secreted factors during hyphal growth. These two *Af* reference strains were chosen as they have different protease activity (30, 31). We previously reported that SARS-CoV2 was able to infect A549 epithelial cells., but showed limited replicative capacity. Increased NLuc signal, indicating viral replication (29), was detected when monolayers were exposed to *Af* culture filtrates from Af293 but not A1160 strains (P < 0.05) **(Figure 2c)**. Interestingly, heightened NLuc activity was detected when A549 cells were co-cultured with THP1 monocytes and Af293 culture filtrates **(Supplementary Figure 2b)**. Increased pathogenicity of *Af* Af293 in our model was further confirmed by increased fungal burdens and stronger suppression of IFNβ release in the presence of Poly(I:C) (P < 0.05) **(Supplementary Figure 2a**,**c)**. Altogether these results suggest that *Af* facilitates viral pathogenesis in a strain dependent manner.

**Figure 2:**
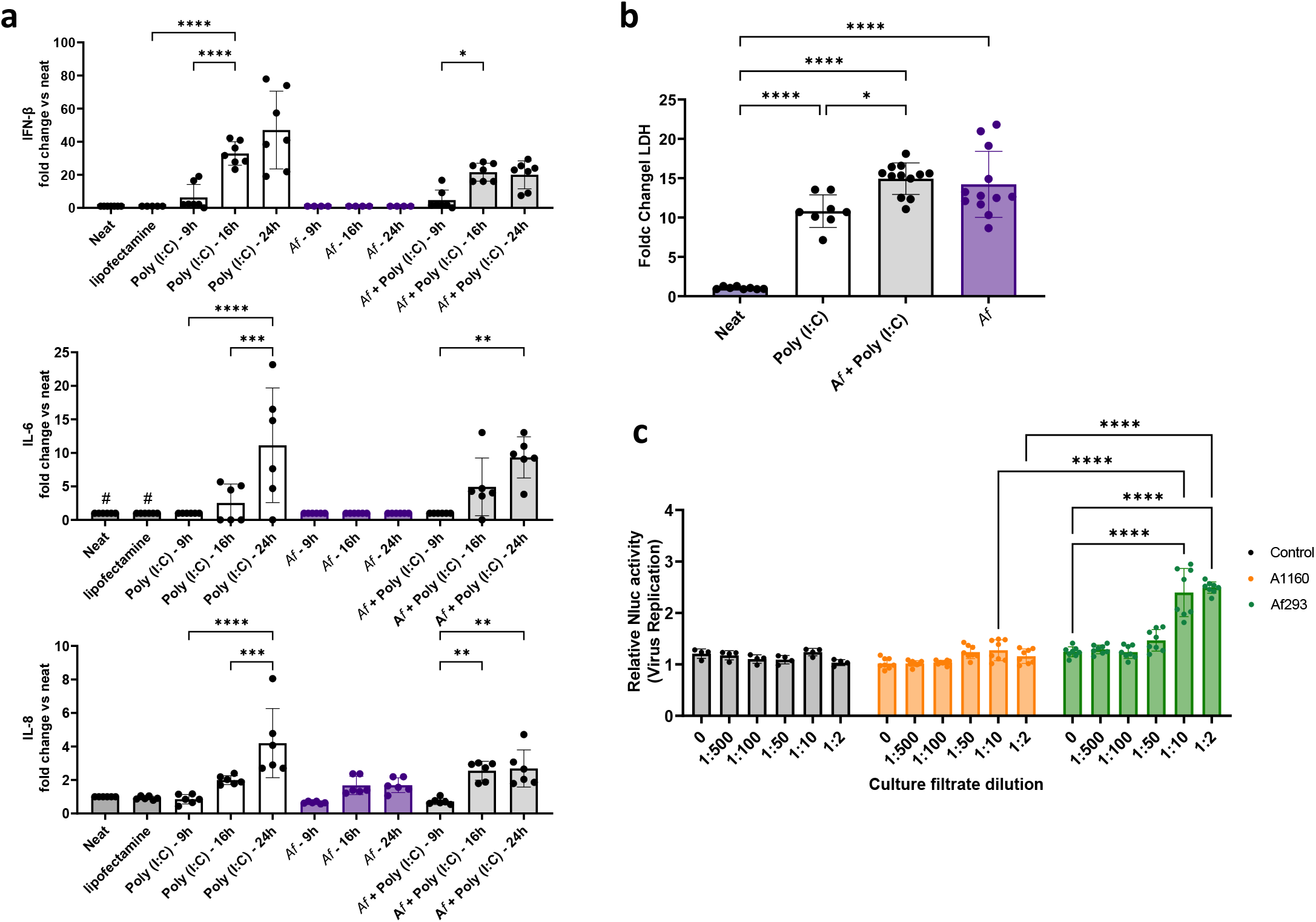
*A. fumigatus (Af)* prevents AECs antiviral responses. (a) AECs cytokine release in the presence of combined *Af* and Poly(I:C) exposure. (b) Viral and fungal exposure does not impact AEC damage compared to single exposure. (c) *Af* enhances SARS-CoV2 infection of A549 cells in a strain-dependent manner. Data in graphs are represented as mean ± standard deviation of biological and technical duplicates/triplicates for experiments with Poly(I:C). For NLuc activity experiments readings for 8 wells were integrated over 200 ms with 4 replicate measurements per well. Mean differences were compared using either one-way or two-way ANOVA with multiple comparison test. \*\*\*\**P* ≤ 0.0001, *** *P* ≤ 0.001, ** *P* ≤ 0.01, and * *P* ≤ 0.05.

Combined infections by fungi and bacteria or viruses are on the rise but the mechanistic understanding about how infection by one pathogen predisposes a host to coinfection or superinfection by another pathogen it is largely unknown. In two independent studies using animal models of disease, it has been shown that *Af* exacerbates pulmonary inflammation during *M. abcessus* and Influenza lung coinfections (32, 33). However, *in vitro* studies have shown that both agonistic and antagonistic interactions might occur between *Af* and pulmonary bacterial pathogens such as *Pseudomonas aeruginosa* in the cystic fibrosis lung [for review (34). Recent transcriptomic data on coinfection of dendritic cells with *Af* and cytomegalovirus (35) or AECs with *A. niger* and SARS-CoV2 (36) suggest that coinfection increases co-pathogenicity. Similarly, it has been reported that antiviral responses of macrophages driven by HIV or Influenza increase *Cryptococcus neoformans* macrophage escape (37, 38). Our findings shed light on the nature of these interactions and suggest that during *Aspergillus*-viral coinfection there is a dual inhibition of antifungal and antiviral AECs responses. The characterisation of what specific fungal factors are master regulators of pathogenesis during coinfection will aid the development of novel therapeutic strategies to prevent and treat these emerging diseases.

### EXPERIMENTAL PROCEDURES

#### Epithelial cell culture

The human pulmonary carcinoma epithelial cell line A549 (ATCC CCL-185) was maintained in supplemented DMEM (sDMEM, 10% foetal bovine serum (FBS), 1% penicillin/streptomycin cocktail). Primary human AECs (Lonza, CC-2547, TAN 40274, 36659 and 40273) were maintained in Small Airway Epithelial Cell growth medium (SAEC, Promocell). Calu-3 bronchial adenocarcinoma cells (ATCC ® HTB-55™) were cultured in supplemented DMEM-F12 (1X) (Thermo-Fisher Scientific). THP-1 (ATCC TIB-202™) monocytes were cultured in supplemented RPMI and added in 1:1 ratio to AECs for co-culture experiments.

All cells were maintained at 37°C, 5% CO_2_ and monolayers used when at ≥ 90% confluence.

#### Fungal strains

The *Af* strains used were A1160, Af293, A1160^+/tdT^ and A1160^+/mSca^ (23), which were cultured and prepared as previously described. For infections, a 10^6^ spores/ml suspension was prepared in sDMEM (for ICF) or Fluorobright sDMEM (Sigma-Aldrich) (for microscopy). The inoculum was verified performing viable counts of serial dilutions on *Aspergillus* complete media (ACM) in duplicate. After the replacement of the culture media with fresh sDMEM (for IFC) or Fluorobright sDMEM (for microscopy), AECs were infected with a final concentration of 10^5^ spore/ml and incubated at 37°C, 5% CO_2_ for the indicated hours in the presence or absence of Poly(I:C). Culture filtrates of *Af* A1160 and Af293 were prepared as described in xx.

#### Viral challenge

Viral challenge was mimicked using Poly (I:C) and, for ICF and microscopy experiments, Poly (I:C)-FITC (Invitrogen). Efficient intracellular delivery of 0.5 μg/ml of Poly (I:C) was achieved by transfection with Lipofectamine 2000 (LP) (Thermofischer) as per manufacturer’s instructions.

#### Effect of viral mimic on antifungal AEC responses

The detailed protocol for single-cell analyses of fungal uptake in cultured AECs using IFC is described in (23, 27). Differently from the protocol cited, due to the use of Poly (I:C)-FITC in our assay, host apoptosis was in this case detected using Annexin V-PE-Cy5 (BioVision) instead of Annexin V-FITC (BioVision). Equivalent quantities were used in the staining mix and fluorescence for this dye was acquired in Channel 5 of the IFC. Live-cell imaging and analyses of fungal uptake and intracellular killing were performed as described in (13) at 6 and 16 h post-infection. Cytokine release (IL-8, IL-6, IFNβ) and LDH release were measured in cell culture supernatants using respectively the DuoSet ELISA kit (R&D) and the CytoTox 96® Non-Radioactive Cytotoxicity Assay (Promega) according to manufacturer’s instructions. *Af* viability in the presence or absence of Poly(I:C) was determined by performing viable counts of serial dilutions after 8h AEC infection.

#### SARS-CoV2–*Af* coinfection

Confluent monolayers of A549 epithelial cells were simultaneously exposed to indicated dilutions of *Af* culture filtrates and a nano-luciferase tagged version of the SARS-CoV2 (SARS-CoV-2-ΔOrf7a-NLuc) as reported in (29). NLuc activity was measured at 72 h post-infection in the presence of 0.5 μL of coelenterazine as previously reported (29). Statistical analyses were done using Prism GraphPad v9 (La Joya, CA).

## Acknowledgements

SG is co-funded by the NIHR BRC Manchester, Fungal Infection Trust (in collaboration with MB), Peel Trust and MAHSC (in collaboration with AP). This work was also supported by a grant to MB from the Medical Research Council (MRC New Investigator Research Grant MR/V031287/1). KE is supported by a NC3Rs training studentship (NC/T001798/1).The authors thank Dr Jennifer Cavet for providing assistance with working at containing level 3 and use of the facilities and Shaunessey Crozier for technical support. The SARS-CoV2 work was done under the approvals of Prof Kadler from the COVID-19 Rapid Response Group, the R3G Research Operations Group and the R3G Executive Committee at the University of Manchester.

## FIGURE LEGEND

**Supplementary Figure 1:** Lipofectamine (LP) neither affect **(a)** *Af* uptake by AECs nor **(b)** FITC-Poly(I:C) intensity. Data in graphs are represented as mean ± standard deviation of biological and technical triplicates. Mean differences were analysed using using one-way ANOVAs and Tukey’s multiple comparison test.. * *P* ≤ 0.05

**Supplementary Figure 2: *A. fumigatus (Af)* prevents AECs antiviral responses in a strain-dependent manner. (a)** Increased Af burdens in AECs in the presence of Poly(I:C). (b) Af293 increases NLuc activity of A549 AECs co-cultured with THP1-monocytes (c) Comparative analyses of Poly(I:C) driven IFNb release by Af culture filtrates. Data in graphs are represented as mean ± standard deviation of biological and technical duplicates/triplicates for experiments with Poly(I:C). For NLuc activity experiments readings for 8 wells were integrated over 200 ms with 4 replicate measurements per well. Mean differences were compared using either one-way or two-way ANOVA with multiple comparison test. \*\*\*\**P* ≤ 0.0001, *** *P* ≤ 0.001, ** *P* ≤ 0.01, and * *P* ≤ 0.05

