## Supplementary figures and images for "Mutual inhibition of airway epithelial responses supports viral and fungal co-pathogenesis during coinfection"

### Supplementary Figures

#### Supplementary Figure 1

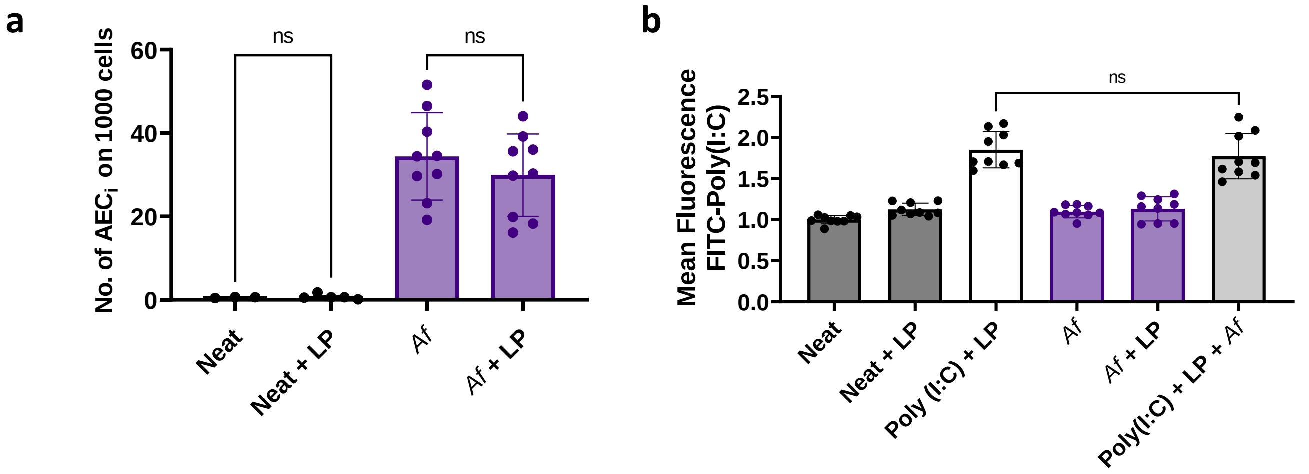

#### Supplementary Figure 2

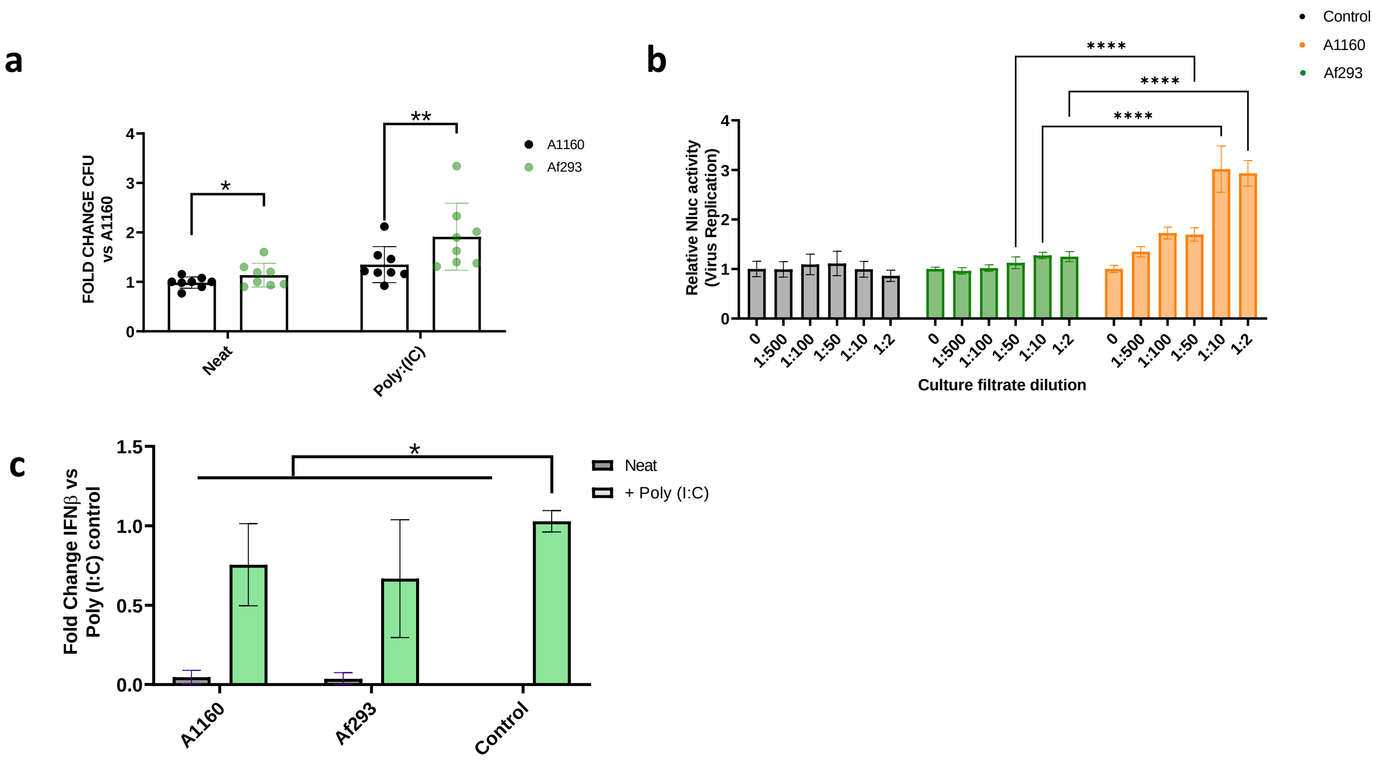
